# Reward positivity biases interval production in a continuous timing task

**DOI:** 10.1101/2023.07.06.548049

**Authors:** Yan Yan, Laurence Hunt, Cameron Hassall

**Affiliations:** Department of Psychiatry, University of Oxford; Department of Psychology, Stanford University; Department of Experimental Psychology, University of Oxford; Department of Psychology, MacEwan University

**Keywords:** Timing, feedback, continuous paradigm, event-related potential (ERP), reward positivity (RewP)

## Abstract

The neural circuits of reward processing and interval timing (including perception and production) are functionally intertwined, suggesting that it might be possible for momentary reward processing to influence subsequent timing behavior. Previous animal and human studies have mainly focused on the effect of reward on interval perception, whereas its impact on interval production is less clear. In this study, we examined whether feedback, as an example of performance-contingent reward, biases interval production. We recorded EEG from 20 participants while they engaged in a continuous drumming task with different realistic tempos (1728 trials per participant). Participants received color-coded feedback after each beat about whether they were correct (on time) or incorrect (early or late). Regression-based EEG analysis was used to unmix the rapid occurrence of a feedback response called the reward positivity (RewP), which is traditionally observed in more slow-paced tasks. Using linear mixed modelling, we found that RewP amplitude predicted timing behavior for the upcoming beat. This performance-biasing effect of the RewP was interpreted as reflecting the impact of fluctuations in dopaminergic activities on timing, and the necessity of continuous paradigms to make such observations was highlighted.

## 1. Introduction

A cortico-striatal dopaminergic system in the brain underlies reward processing (Smith and Kieval, 2000; Schultz, 2007; Arias-Carrión et al., 2010; Haber and Knutson, 2010; Corlett et al., 2022). Interestingly, the same dopaminergic circuits have been implied in beat-based interval timing, where the interval is encoded relative to the beat (Rao et al., 2001; Matell et al., 2003; Matell and Meck, 2004; Grahn, 2009; Coull et al., 2011; Teki et al., 2012). This overlap in functional circuits for reward and timing raises the possibility that phasic activation of reward circuits during reward processing can lead to temporary changes in timing behavior. Such interaction may be more prominent for sub-second than supra-second interval timing, due to more intense activation of subcortical areas (Wiener et al., 2010; Nani et al., 2019).

Interval timing includes interval perception and interval production (Ivry and Hazeltine, 1995). Past literature has predominantly focused on the effect of reward on interval *perception,* while the effect of reward on interval *production* remains understudied. Animal studies demonstrated that direct manipulation of phasic dopamine signaling alters interval perception. For example, Soares et al. (2016) reported that spontaneous fluctuations in phasic DA signaling lead to mice perceiving the same interval as shorter, and optogenetic manipulation is sufficient for reproducing such behavioral pattern. In humans, researchers typically studied the influence of externally administered reward on interval perception. It was shown that intervals associated with a monetary positive prediction error or reward were perceived as longer by human participants (Failing and Theeuwes, 2016; Toren et al., 2020), an effect partially attributable to attention and salience (Berridge and Robinson, 1998; Tse et al., 2010). However, these studies typically manipulated reward independently of performance, whereas in real life, reward is likely contingent on timing performance (Ariely and Zakay, 2001; Petter et al., 2018).

Feedback is an incidence of performance-contingent reward, and positive feedback on performance (e.g., ‘accurate’, ‘hit’ or ‘on time’) is known to elicit a reward response in the brain 200-300 ms post-stimulus (Proudfit, 2015; Cockburn and Holroyd, 2018; Tunison et al., 2019). While several previous studies used fMRI to study the continuous sub-second interval production with visual feedback (e.g., Lutz et al., 2000; Pope et al., 2005), it may be difficult to conduct a trial-by-trial fMRI analysis to capture the transient changes in reward circuit due to constraints on temporal resolution. EEG allows for fine-grained discrimination of such interplay at the level of milliseconds, making it especially suitable for studying the influence of reward on sub-second timing behavior. It has been well-established that reward, relative to non-reward, elicits a positive deflection in the scalp-reported event-related potential (ERP) in the frontocentral electrode sites around 250 to 350 ms after stimulus onset called the Reward Positivity (RewP) (Holroyd and Coles, 2002; Holroyd et al., 2006, 2011; Walsh and Anderson, 2012)^1^. RewP variability is likely due to variability in the reward response rather than the non-reward response (Holroyd et al., 2008). One theory of the RewP highlights its link to the reward prediction error in reinforcement learning (Sutton and Barto, 2018), attributing the RewP to the influence of a phasic ventral striatal DA signal on the anterior cingulate cortex (Holroyd and Coles, 2002; Luu et al., 2003; Carlson et al., 2011). This outcome information is utilized by the anterior cingulate cortex to compute a need-for-control signal, facilitating cognitive control and effort exertion (Shenhav et al., 2013; Vassena et al., 2017). Altogether, the RewP provides a non-invasive, temporally sensitive measure of reward prediction errors on the scalp.

In this study, we asked how reward biases sub-second interval production in a continuous timing paradigm with EEG recording. Participants were instructed to reproduce different drumming patterns at different tempos (fast, medium, and slow) using two keys on a keyboard, and received color-coded feedback (early, on time, or late) on their accuracy after each response. We hypothesized that on-time feedback would elicit a RewP relative to early or late feedback, and examined whether the RewP could be reliably observed in all three tempos. We then hypothesized that trial-to-trial instantaneous fluctuations of RewP amplitude in response to on-time feedback biases subsequent interval production, using a linear mixed model. We reported that RewP was only stably observed in the medium and slow tempos. In the slow tempo, a larger RewP in response to ‘on time’ feedback led to a longer produced interval on the next trial.

## 2. Methods

### 2.1 Participants

21 participants completed the study. One participant (No. 12, female) was excluded due to the trigger cable being partially disconnected. The remaining 20 participants (15 female, 2 left-handed, mean age 25.85 ± 4.53) had normal or corrected-to-normal vision and had no known neurological impairments. All participants gave informed consent and were compensated for their participation and a performance bonus. This study was approved by the Medical Sciences Interdivisional Research Ethics Committee at the University of Oxford (R51132/RE002).

### 2.2 Experimental Task

Participants completed the drumming task along with two other unrelated tasks, the order of which was randomized between participants. Each task took approximately 20 minutes, and the entire study took around 60 minutes. The stimuli were presented using PsychoPy 3.6.6 (Peirce et al., 2019) on a monitor screen with size 59.9 cm (width) × 33.7 (height). In each block of the drumming task, participants listened to a drumming pattern for 24 beats and were asked to reproduce the pattern using the F and J keys on the keyboard (**Figure 1A)**. The response sequence was self-initiated by pressing the first key, and participants were shown color-coded visual feedback for 50 ms after each subsequent key press response, indicating if their response was fast, on time, or slow. The palette is color blind-friendly and the correspondence between color and feedback was counterbalanced across participants. If the participant pressed a wrong button, a red ‘X’ appeared on the screen instead for 50 ms. A participant was ‘on time’ if the produced interval fell into the target interval plus or minus a given margin. The margin had a starting width of 100 ms, and a staircase procedure was used to adjust the size of this margin by steps of 10 ms, so that in every block type, the ‘on time’ feedback type composed approximately 50% of all feedbacks. There were three tempo conditions: fast (150 beats per minute [BPM], target interval 0.4 s) medium (100 BPM, target interval 0.6 s), and slow (60 BPM, target interval 1 s). In each condition, there were two possible drumming patterns ‘aaba/aaba/aaba…’ (commonly referred to as ‘4/4 time’) and ‘aaabaa/aaabaa/aaabaa…’ (commonly referred to as ‘6/8 time’). Both hands were used as starting hand to balance for dominant hand effect. Each participant completed 24 blocks in total (2 patterns × 3 tempos × 2 dominant hands × 2 repetitions). Participants received a bonus for their performance on the task. Twenty of the 21 participants completed 24 blocks (72 trials per block, 1728 trials in total) in total. One participant (No. 1, female) completed a pilot version of the task which included two identical repetitions of the task as performed by other participants, and only the first 1728 trials were included in the analysis.

**Figure 1.**
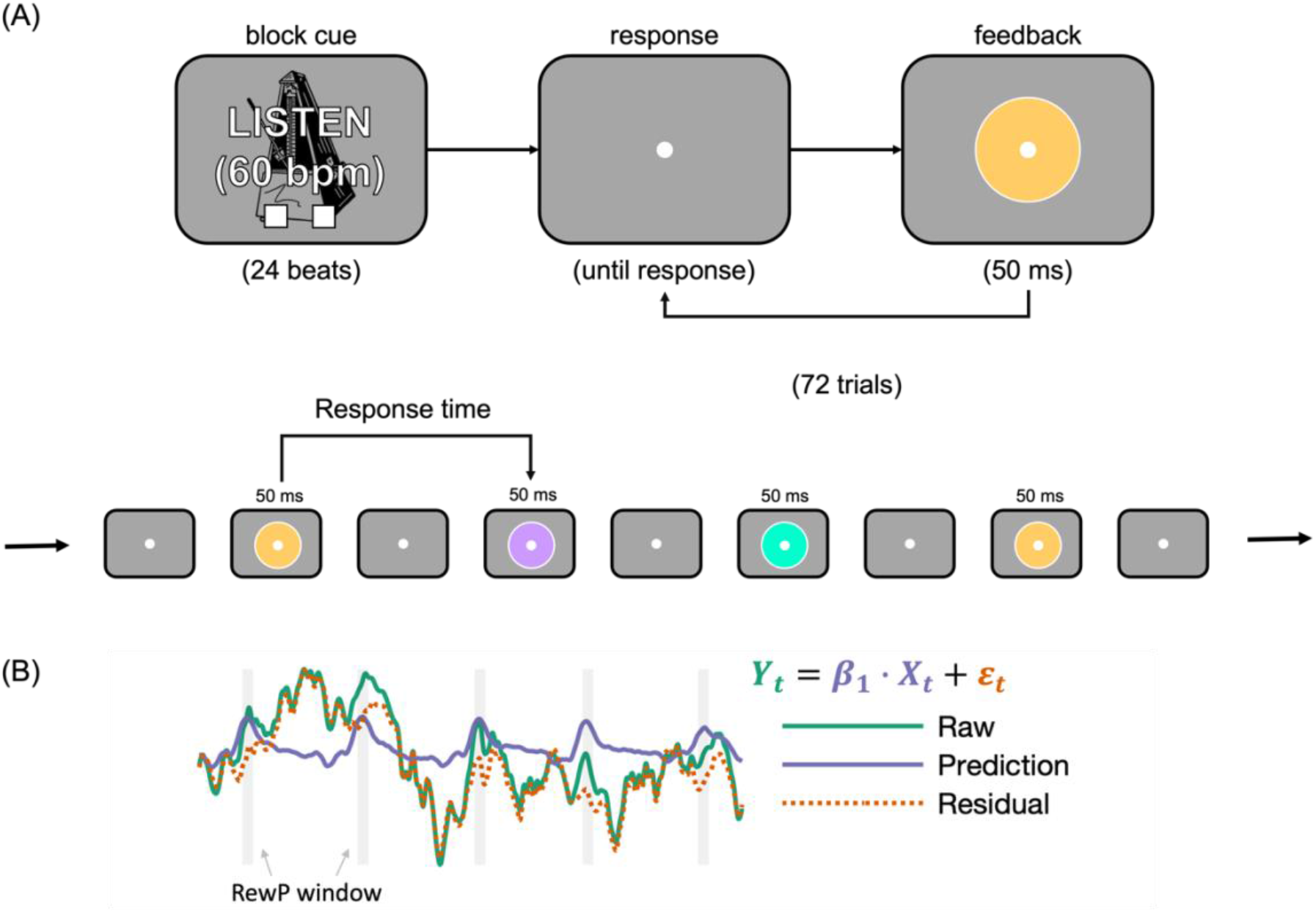
Task design and EEG data analysis. (A) The drumming task. In each block, participants were required to produce a drumming pattern using F and J keys on the keyboard. Each response was provided with color-coded visual feedback indicating if it was early, on time or late. There were three tempos in the experiment: fast, medium, and slow. (B) The calculation of trial-to-trial residual RewP. The residual EEG was calculated by subtracting the predicted EEG amplitude from the regression analysis from the raw EEG amplitude at each time point. A trial’s RewP was the average amplitude of residual EEG in the RewP time window, shown as grey rectangles in the figure.

This drumming task is continuous in the sense that participants self-paced the drumbeats and there was no artificial delay between events, making it relatively naturalistic. Because the timing response immediately occurred 50 ms after the onset of feedback stimulus, signaling the start of the next timing interval, there were no ‘trials’ in the traditional sense. Instead, we defined a ‘trial’ in this paradigm as starting with feedback onset, followed by feedback-relayed neural responses, and until the subsequent button response. Similarly, we defined response time (RT) as each inter-beat interval between two drumbeats generated by key presses.

### 2.3 Software

Preprocessing and analysis of EEG data was conducted in MATLAB R2020a (The MathWorks Inc., 2020), using EEGLAB (Delorme and Makeig, 2004). The results were then analyzed in RStudio (version 4.0.2, 2020-06-22). Linear mixed models were conducted using the R package *lme4* version 1.1-31 (Bates et al., 2014) and *lmerTest* version 3.1-3 (Kuznetsova et al., 2017), and effect size estimates were acquired using the R package *effectsize* version 0.8.2 (Ben-Shachar et al., 2020) and package *EMAtools* version 0.1.4.

### 2.4 Behavioral Analysis

#### 2.4.1 Response Time and its adjustment

Response time (RT) was defined as the time to press the button after feedback onset. RT adjustment was calculated as the difference between RT in the current trial and in the last trial. A positive value suggests increasing RT, and a negative value suggests decreasing RT. To analyze timing performance, we first removed trials where participants pressed the wrong button, and excluded outliers of RT and RT adjustment in the top and bottom 1%. We conducted a one-sample t-test comparing participants’ average response time in each tempo to the target interval and reported Cohen’s d (Cohen, 2013). We conducted a two-way ANOVA (3 tempos × 3 feedback types) on participants’ average RT adjustment, and reported the effect size partial eta squared (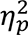).

#### 2.4.2 Hierarchical timing and chunking

During execution of movement sequences, participants tend to group consecutive movements together, and organize them in a hierarchical manner (Rosenbaum et al., 1984; Verwey and Dronkert, 1996; Verwey and Eikelboom, 2003; Sternberg et al., 2018). The chunking literature suggests that participants’ movement is more fluent and efficient within a chunk than when they switch between chunks (Verwey and Dronkert, 1996; Ramkumar et al., 2016). Importantly, chunking has been linked to dopaminergic functioning in animals and humans (Tremblay et al., 2009, 2010), and chunking may lead to phasic DA signaling as a function of the relative location of the current beat in a chunk. In this task, participants were explicitly instructed to drum at a specific pattern, so we used the current beat’s location in the specific pattern as a proxy for chunking. We analyzed the effect of chunking on RT and its adjustment, and accounted for chunking in the linear mixed models (for details, see **section 2.5.5**).

#### 2.4.3 Regression to local mean

One possible concern with our RT adjustment measure is that it could partially reflect regression to the mean. A short RT could be followed by an apparent ‘adjustment’ in the right direction, simply because the next response is more likely to be closer to the mean RT (Jazayeri and Shadlen, 2010). To address this issue, we conducted a simulation by drawing response time (1728 draws per tempo) from a Gaussian distribution specified by the observed mean and standard deviation, and derived apparent ‘RT adjustment’ as the difference between consecutive RT draws. If the observed RT adjustment following different feedbacks in this task is not different from the simulated null distribution, then we may conclude that the apparent RT adjustment observed in the study likely arose only from regression to the mean. Moreover, if RT adjustment is only due to regression to the mean, we should not observe any effect of neural processing of feedback such as RewP.

Regression to the mean suggests that a larger deviation from the mean leads to larger subsequent adjustment in the opposite direction, resulting in a negative association between the two values. Moreover, it is likely that participants’ performance drift over time, shifting the distribution from which the current RT is drawn. Therefore, regression to the mean ought to be quantified relative to the *local* mean (e.g., the recent trials), but not the grand mean (e.g., the mean RT in the current block or the target interval). This deviation-from-mean parameter was quantified as the difference between the RT on this trial and the local mean RT, which was the rolling mean averaged across the previous 10 trials. For the first 10 trials in a block where this rolling mean cannot be calculated, the deviation was calculated as the difference between RT and the target interval (0.4, 0.6, or 1 s). A positive value of this deviation variable suggests temporary slowing on this trial compared to recent history, and a negative value suggesting temporary speeding. We demonstrated using simulation that when apparent RT adjustment solely arises from regression to the mean in a Gaussian distribution, this adjustment is negatively correlated with deviation from local mean (**Supplementary Information Figure S1.3 A-B**). Therefore, in the linear mixed modeling, we added deviation from the mean as a covariate to partial out the effect of regression to the mean on RT adjustment. Varying the time window for calculating the rolling mean, or excluding the first trials where the rolling mean could not be calculated did not alter the main conclusions from the linear mixed model.

### 2.5 EEG Analysis

#### 2.5.1 EEG Recording

32 channel EEG was recorded at 1000 Hz with an actiCHamp Plus amplifier (Brain Products, GmbH, Gilching, Germany) using BrainVision Recorder (Version 1.23.0001, Brain Products, GmbH, Gilching, Germany). The EEG recording was referenced to Fz online. 30 of the electrodes were arranged according to the international 10-20 system, and two additional electrodes were placed on the left and right mastoids.

#### 2.5.2 Pre-processing

The EEG was pre-processed in MATLAB R2020a (Mathworks, Natick, USA) using EEGLAB (Delorme and Makeig, 2004). EEG data was down-sampled to 250 Hz, filtered by a 0.1-30 Hz band pass filter and a 50 Hz notch filter, and re-referenced to the linked mastoids. Ocular artefacts were identified and removed from the continuous data by running an independent component analysis and then the *iclabel* function.

#### 2.5.3 Extraction of Regression-based Event-Related Potential

In this continuous task, each behavioral response was immediately followed by visual feedback, and the interval timing for the next beat immediately ensued without an inter-trial interval. Due to component overlap, this rapid design poses challenges to the traditional event-related approach to EEG analysis. We used a regression-based ERP (rERP) analysis method to extract waveforms from the overlapping signals using the Unfold toolbox in MATLAB (Smith and Kutas, 2015; Ehinger and Dimigen, 2019). We detected artifacts in the continuous EEG with a 150 μV threshold using the *basicrap* function from ERPLAB toolbox with 2000 ms window and 1000 ms step size (Lopez-Calderon and Luck, 2014). For each participant, we constructed a design matrix consisting of stick functions spanning -1500 to 1500 ms around the onset of visual feedback, for each feedback type and tempo, respectively. EEG sample and design matrix rows corresponding to artefacts were removed before solving the equation. We also conducted a traditional EEG analysis for comparison (**Supplementary Information S2**).2.5.4 RewP Amplitude Quantification To identify the scalp location of the RewP, we used the ‘collapsed localizer’ approach (Luck and Gaspelin, 2017), combining across tempos and incorrect feedback types (early or late) to form a single correct waveform and a single incorrect waveform for each electrode. We located the electrode (FCz) at which the RewP amplitude (collapsed correct minus collapsed incorrect) was maximal. RewP time window was selected as 240-340 ms according to a previous meta-analysis (Sambrook and Goslin, 2015). RewP was calculated as the difference wave between correct and incorrect feedback, and the amplitude is quantified as the mean amplitude in the RewP time window. We conducted one-sample t-test comparing participants’ average amplitude to 0 and reported Cohen’s d (Cohen, 2013). We then conducted a two-way within-subject ANOVA (3 tempos × 2 feedback contrasts) on participants’ average RewP amplitude, reporting the partial eta-squared. The average waveform and its topography were plotted by averaging across all participants for three tempos respectively.

#### 2.5.5 Trial-by-trial analysis

After confirming the existence of RewP and localizing it to the electrode FCz, we asked the question whether the neural activity at this electrode site induced by reward (i.e., on time feedback) biases subsequent timing. We used the trial-by-trial EEG amplitude during on-time trials in the RewP time window as a proxy to reward-induced dopaminergic fluctuations, and use this amplitude to predict participants’ behavioral adjustment in the next trial. We focused on the on-time trials for two reasons. First, there is evidence that RewP variability depends on the reward response but not the non-reward response (Holroyd et al., 2008; Proudfit, 2015). Second, reward feedback in this task ought to be unconfounded by the directional behavioral adjustments we would expect for ‘early’ and ‘late’ feedback (**SI table S1.1**).

To derive the trial-by-trial RewP amplitude, we extracted a common feedback component for each participant’s individual condition (fast, medium, slow), and feedback type (early, late, on time), using the *r*ERPs acquired from the regression-based analysis above. The trial-to-trial ‘residual RewP’ was computed as the difference between the current trial’s EEG amplitude and the predicted amplitude from the regression model, averaged within the pre-specified RewP window (240-340 ms) at electrode FCz (**Figure 1B**). As a comparison, we also derived the trial-to-trial residual EEG from the traditional ERP approach (**Supplementary Information S2)**. This experiment does not contain explicit practice trials; the first three blocks were considered as practice blocks and excluded from the analysis. We further truncated the top and bottom 1% of RT for each tempo, 1% of all RT adjustment, regression to the mean, and residual RewP amplitude from the dataset, removing 7.0% of all trials. For sensitivity analysis, we varied the number of blocks counted as practice blocks, and the percentage of outliers (see **Results 3.3**). Before exclusion, the range of RT was 0.001 to 25.573 seconds, the range of RT adjustment was -24.995 to 24.949 seconds, the range of deviation from the mean was -4.004 to 22.118 seconds, and the range of trial-to-trial EEG was -76.786 to 66.371 μV; after exclusion, the range of RT was 0.305 to 1.222 seconds, the range of RT adjustment was -0.290 to 0.283 seconds, the range of deviation from the mean was -0.419 to 0.391 seconds, and the range of trial-to-trial EEG was -28.593 to 28.777 μV. Linear mixed models were constructed using tempo, feedback type, residual RewP and their interactions to predict the RT adjustment (signed; positive value indicates slowing) on the next trial, while controlling for chunking and regression to the mean. We focused on the slow tempo, where the most prominent RewP was observed (**Section 3.2**), and on the on time feedback type, because this is where the hypothesized fluctuations in phasic DA signaling occurs. We constructed the following linear mixed model using the R function *lmer()* with random intercept for each participant:

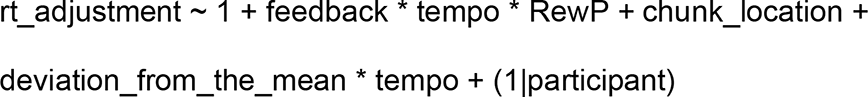

Here, chunk location denotes the location of the current interval in the drumming pattern. The main effect of RewP on RT adjustment following on time feedback within each tempo was acquired by relevelling the model to different tempos and re-running the model. Cohen’s d was reported for all linear regressions. Finally, we fit a linear mixed model with random slopes and intercept for every participant:

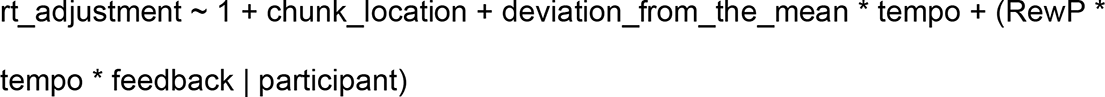

We extracted the slope coefficients for each participant respectively and tested whether they are systematically greater than 0 in a one-sample t-test.

## 3. Result

### 3.1 Systematic bias in interval production

Participants were relatively accurate in reproducing the target intervals (**Figure 2A-B**). One-sample t-tests on participant’s average RT indicated that participants were significantly faster than the target interval for the medium tempo (all RT below in seconds; Mean difference = 0.570 s, SD = 0.020, t_(1,19)_ = -6.867, p < .001, Cohen’s d = -1.54) and the slow tempo (Mean difference = 0.933 s, SD = 0.037, t_(1,19)_ = -7.884, p < .001, Cohen’s d = -1.76), but not for the fast tempo (Mean difference = 0.402 s, SD = 0.014, t_(1,19)_ = 0.661, p = .516, Cohen’s d = 0.15). Consistent with the systematic bias in RT, participants also received asymmetric proportion of ‘early’ and ‘late’ feedbacks, while the proportion of on time feedback was approximately 50% (**SI Table S1.1**). Chunking was observed for both drumming patterns; participants’ RTs were faster when they were within a chunk, than when they moved to another chunk or switched hands (**SI Figure S1.3C**).

**Figure 2.**
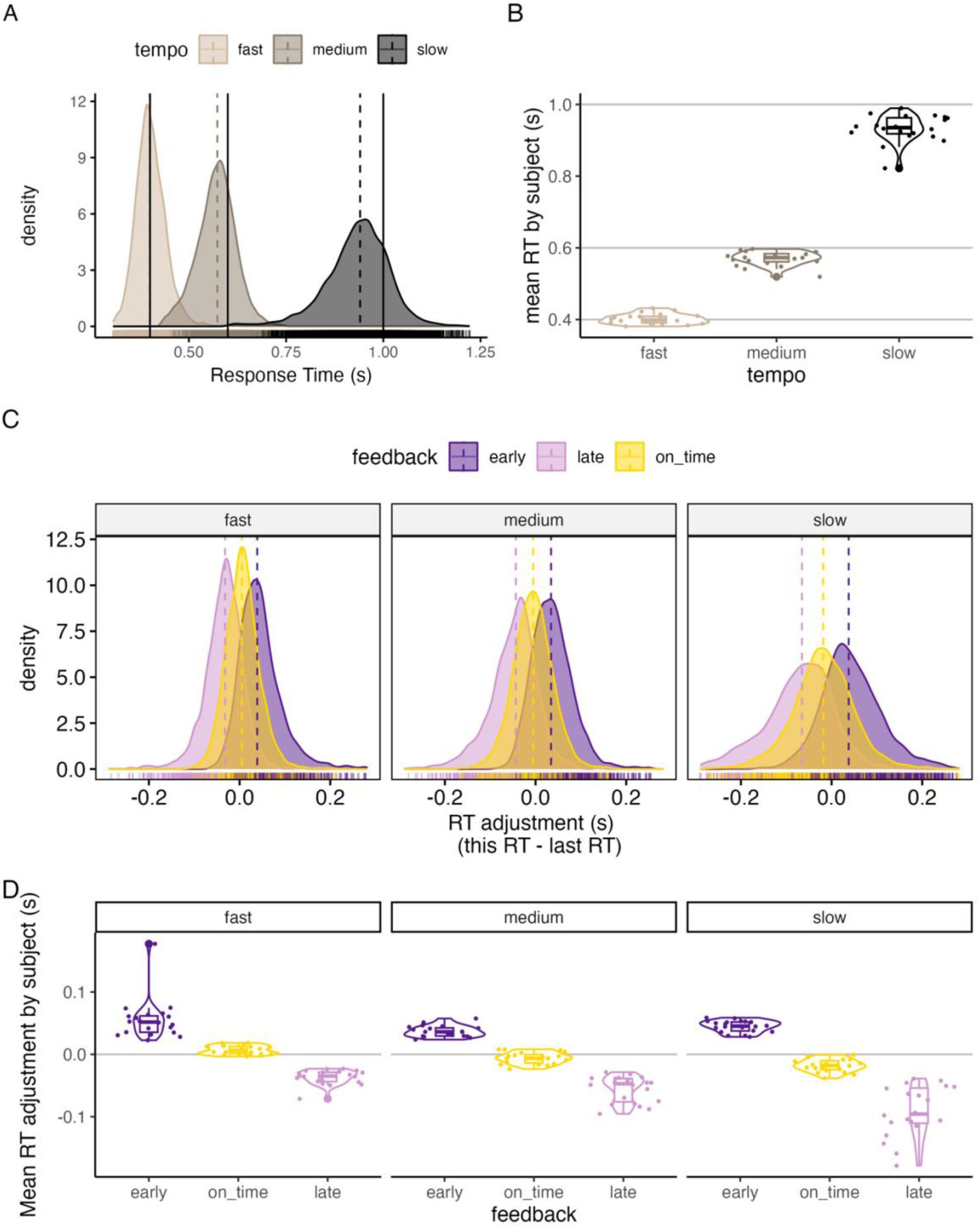
Participants relatively accurately reproduced different drumming patterns and adjusted their behavior following different feedbacks. (A) The distribution density of RT in each tempo, suggesting speeding in the medium and the slow tempo. The dotted line and the solid line indicated the population mean and the target interval, respectively. (B) The distribution of participants’ average RT in each tempo. Each point represents one participant. The horizontal lines represent the target interval. (C) The distribution of RT adjustment in each tempo. The dotted lines showed the population mean. (C) The distribution of RT adjustment by feedback type in each tempo.

Two-way within-subject ANOVA (3 tempos × 3 feedback types) on participants’ average RT adjustment suggested that RT adjustment significantly differed by feedback type (F_(2,38)_ = 205.651, p < .001, 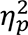 = 0.92) and tempo (F_(2,38)_ = 28.213, p < .001, 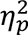 = 0.60). Importantly, the significant main effect of feedback type confirms that participants adjusted their behavior according to feedback, speeding up upon receiving ‘late’ feedback and slowing down upon receiving ‘early’ feedback (**Figure 2C**). There was also a significant interaction between feedback and tempo (F_(4,76)_ = 21.280, p < .001, 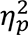 = 0.53). Pairwise comparisons with Bonferroni adjustment further suggested that participants’ RT adjustment following ‘on time’ feedback was significantly more positive (suggesting slowing) for fast tempo than slow tempo (mean difference = 0.025 s, t-ratio = 3.957, p_adjust_ < .001, Cohen’s d = 1.83). The biasing effect of tempo and feedback on RT was systematic across participants, although there were individual differences in their mean RT and RT adjustments (**Figure 2D**).

### 3.2 RewP was observed only for the medium and the slow tempos

We derived rERPs for each tempo and feedback type (**Figure 3A-C**). The slower the tempo, the larger the Reward Positivity observed (**Figure 3D-F**). In the slow tempo, but not the medium and fast tempo, a clear frontocentral gradient of scalp RewP amplitudes emerged that peaked at FCz (**Figure 3G-I**). Two-way within-subject ANOVA (3 tempos × 2 feedback contrasts) on the mean RewP amplitude for each participant suggested a significant main effect of tempo (F_(2,38)_ = 7.560, p = .002, 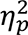 = 0.285), but not feedback type (F_(1,19)_ = 0.044, p = .837, 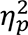 = 0.002). There was no interaction between tempo and feedback type (F_(2,38)_ = 0.270, p = .765, 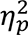 = 0.014). Pairwise t-tests with Bonferroni corrections suggested that the RewP amplitude in the slow tempo was significantly larger than in the medium tempo (Mean difference = 0.931, t_(1,39)_ = 2.474, adjusted p = .018, Cohen’s d = 0.39) and in the fast tempo (Mean difference = 1.768, t_(1,39)_ = 3.509, adjusted p = .001, Cohen’s d = 0.55); the RewP amplitude in the medium tempo was significantly larger than that in the fast tempo (Mean difference = 0.837, t_(1,39)_ = 2.043, adjusted p = 0.048, Cohen’s d = 0.32). The 95% confidence interval of the RewP amplitude in the fast tempo included zero, and the 95% confidence interval of the amplitude from medium and slow tempo were greater than 0, suggesting that RewP could be reliably observed in the medium and slow tempo (**Table 1**).

**Figure 3.**
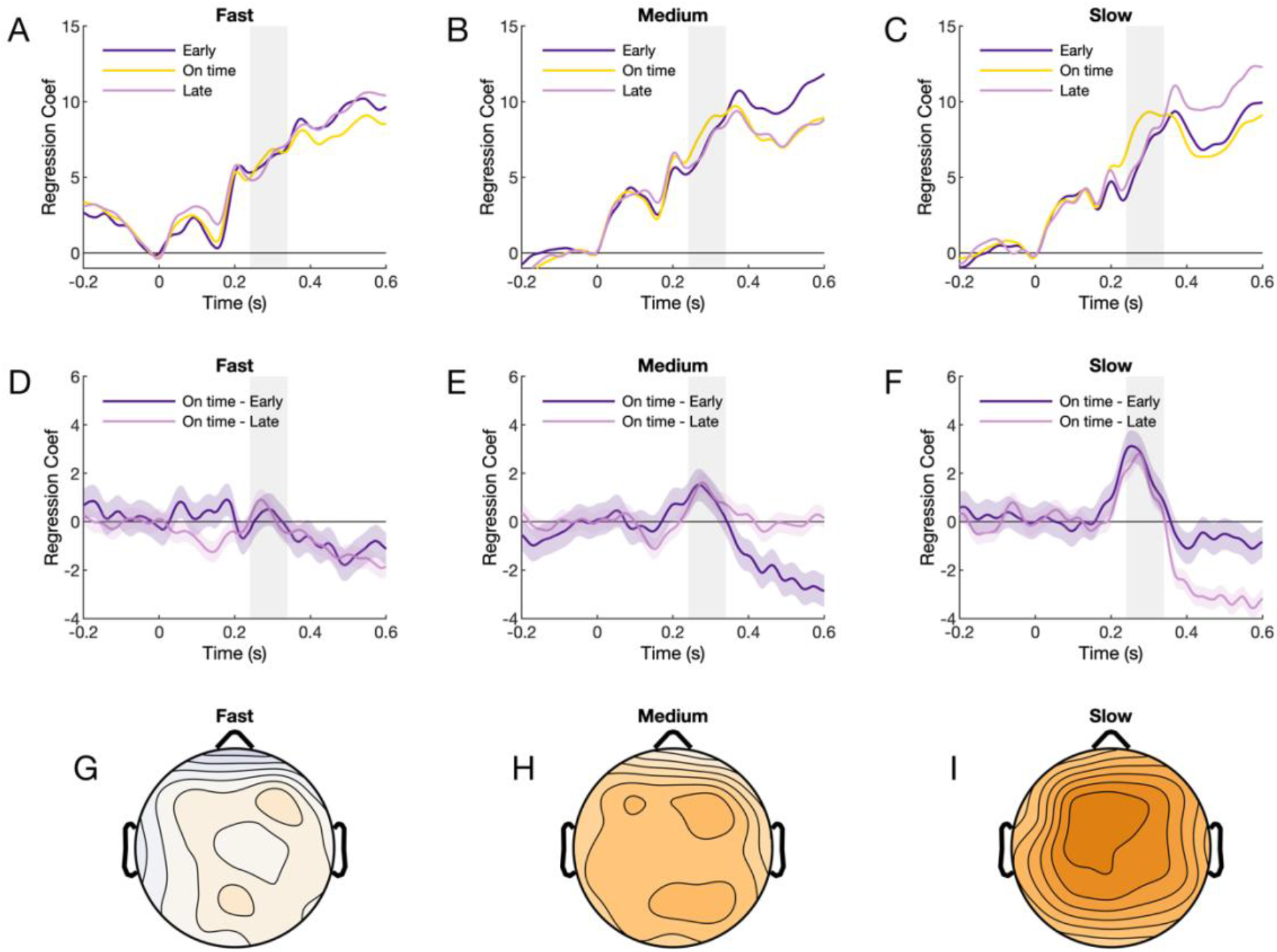
Reward Positivity in the fast, medium and slow tempo. (A-C) Regression-based ERP by tempo and feedback type. The RewP time window as reported in Sambrook and Goslin (2015), was highlighted in grey (240-340 ms). (D-F) The RewP wave was calculated as the contrast between correct and incorrect feedback type. RewP amplitude increased as a function of target interval. (G-H) The topography of RewP, averaged between early and late feedback type. In the slow tempo, the peak amplitude was located at electrode FCz.

**Table 1.**
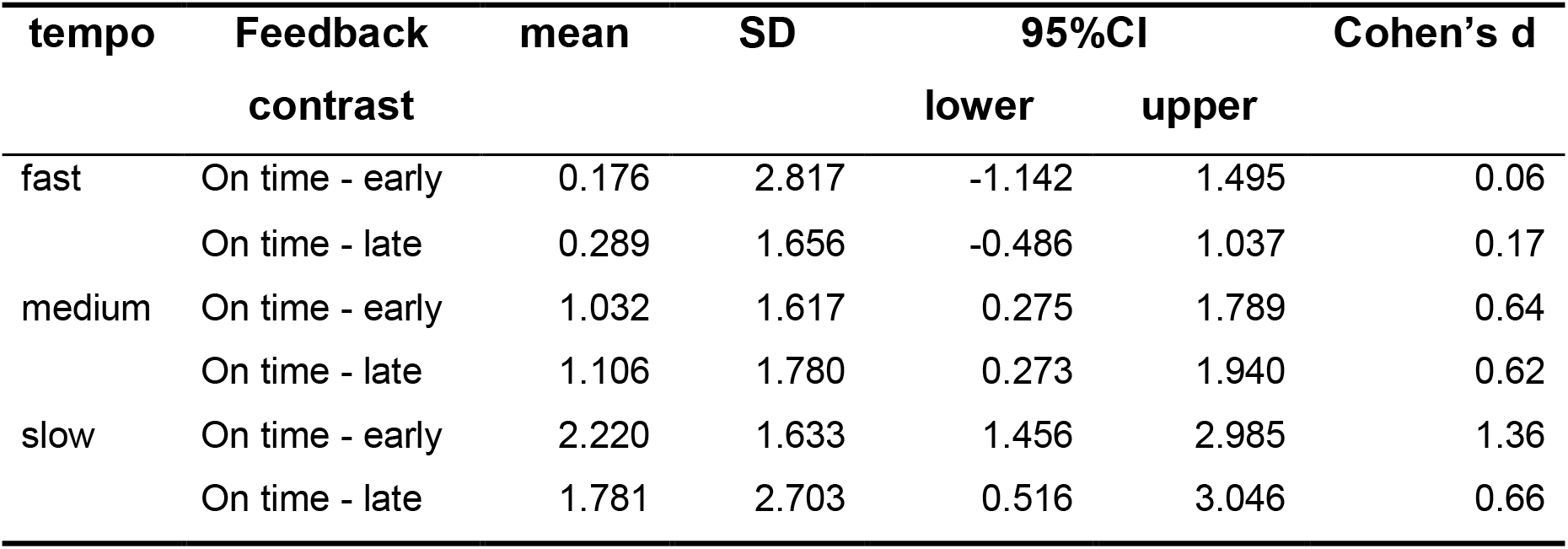
Summary statistics of participants’ mean RewP amplitude for each tempo and feedback type.

| tempo | Feedback | mean | SD | 95%CI |  | Cohen's d |
| --- | --- | --- | --- | --- | --- | --- |
|  | contrast |  |  | lower | upper |  |
| fast | On time - early | 0.176 | 2.817 | -1.142 | 1.495 | 0.06 |
|  | On time - late | 0.289 | 1.656 | -0.486 | 1.037 | 0.17 |
| medium | On time - early | 1.032 | 1.617 | 0.275 | 1.789 | 0.64 |
|  | On time - late | 1.106 | 1.780 | 0.273 | 1.940 | 0.62 |
| slow | On time - early | 2.220 | 1.633 | 1.456 | 2.985 | 1.36 |
|  | On time - late | 1.781 | 2.703 | 0.516 | 3.046 | 0.66 |

### 3.3 Trial-to-trial fluctuation in RewP amplitude biases timing in peri-second range

We examined the autocorrelation of RT adjustment and RewP using linear mixed models with random intercept for each participant. Last trial’s RT adjustment was negatively associated with RT adjustment on this trial, such that slowing on the last trial predicted speeding up on this trial, and vice versa (regression coefficient *B* = - 0.257, t = -61.281, p < .001, Cohen’s d = -0.749), suggesting regression to the mean. Last trial’s RewP was not associated with this trial’s RewP, *B* = -0.0016, t = 0.316, p = 0.752, Cohen’s d = 0.004, suggesting that there are negligible baseline fluctuations in RewP across trials.

We divided the RewP amplitudes into 10 bins with 10% of trial data in each bin, and plotted the mean and standard error of RT adjustment for this RewP bin. Visual inspection revealed a linear association between RewP amplitude and RT adjustment (**Figure 4A**). Next, we fitted a linear mixed model using the current trial’s residual RewP (trial EEG subtracted by average EEG waveform) to predict RT adjustment on the next trial. The model converged successfully. Feedback type, tempo, regression to the mean, and chunk locations significantly predicted RT adjustment. In the slow tempo, trial-to-trial fluctuations in RewP in response to on time feedback predicted RT adjustment on the next trial, such that larger (more positive) RewP led to a slowing of RT compared to the last trial (*B* = 2.9×10^-4^, t = 4.341, p < .001); the effect size of the biasing effect of RewP was modest, Cohen’s d = 0.053 (**Figure 4B**). Such timing-biasing effect of RewP fluctuations was not observed for the fast (*B* = 1.2×10^-4^, t = 1.892, p = 0.058, Cohen’s d = 0.023) or the medium tempo (*B* = 0×10^-4^, t = -0.002, p = 0.999, Cohen’s d < .001); RewP fluctuations did not bias timing following early or late feedback (|t| < 1.390, p > 0.164). The effect of tempo and chunk location on RT adjustment, and estimates of the entire model, are shown in **Supplementary Information (Table S1.1)**. The regression coefficient *B* of RewP was still significant when not including chunk location and regression to the mean by tempos as the covariates, B = 2.7×10^-4^, t = 3.703, p < .001, Cohen’s d = 0.045.

**Figure 4.**
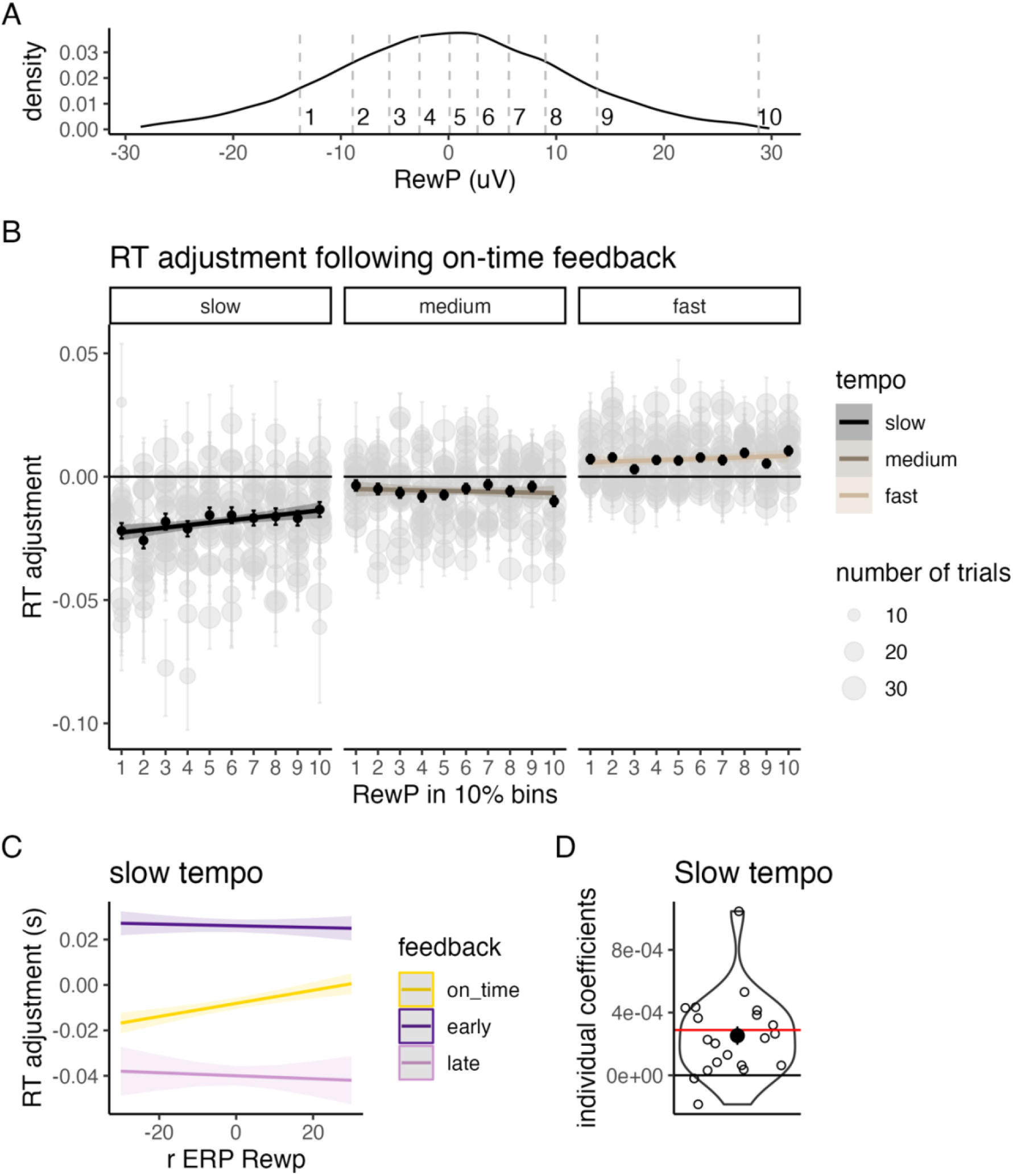
Reward response predicted timing behavior. (A) For visualizing RT adjustment as a function of RewP, all RewPs were divided into 10 bins of 10%. (B) Size of RT adjustment as a function of the RewP amplitude in response to on time feedback. Each grey point represented one participant’s data in this bin, with larger point size indicating more trials. The grey-colored error bars shown below indicated standard error of the subset of data represented by the grey point. The black points and error bars represent the group-level mean and standard error within each RewP bin. (C) The predicted values of RT adjustment in the slow tempo from the linear mixed model. (D) Individual regression coefficients *B* from a linear mixed model with random slopes and random intercept, in the slow tempo. Each open dot represents the value of one participant, and red line denotes the coefficient value when the slope is fixed across participants.

To confirm that the finding that trial-to-trial RewP was not limited to the present exclusion criterion, we conducted sensitivity analysis by varying (1) the number of blocks considered as practice blocks, and (2) the percentage of outliers in timing behavior and EEG. When no block was counted as practice block, the regression coefficient of RewP was still significant, B = 2.95×10^-4^, t = 4.431, p < .001, Cohen’s d = 0.051. When only the bottom and top 0.5% of RT, RT adjustment, deviation from the mean, and RewP amplitude were excluded (inclusion rate 95.76%), the regression coefficient was still significant, B = 2.8×10^-4^, t = 3.742, p = .002, Cohen’s d = 0.042.

As shown in **Figure 4C**, when fitting both random slope and random intercept for each participant’s RewP predicting RT adjustment, all but two participants’ slope estimates were larger than 0 (min = -0.18×10^-4^, max = 10.5×10^-4^, Mean = 2.5×10^-4^, SD = 2.6×10^-4^). One-sample t-test suggests that the slopes were significantly larger than 0 (t_(1,19)_ = 4.304, p < .001, Cohen’s d = 0.96).

## Discussion

This study investigated the influence of reward processing on interval production by looking at participants’ Reward Positivity (RewP) in response to rapid feedback while they engaged with a continuous drumming task at different tempos. A continuous timing paradigm was used to gather larger number of trials compared to traditional trial-based paradigms. Trial-to-trial EEG fluctuations in the RewP time window predicted timing adjustment on the next trial, such that a larger (more positive) RewP amplitude relative to the mean waveform forecasted longer produced interval on the next trial. This study demonstrated the plausibility of using a rapid paradigm to acquire the RewP, and showed that fluctuations in the RewP are associated with variations in interval production.

Considering previous studies that linked the RewP to a striatal reward prediction error relayed to anterior cingulate cortex (Holroyd and Coles, 2002; Holroyd and Yeung, 2012), our findings could be interpreted as a slowing effect of reward-related phasic DA signaling on interval production. Previous studies have reported that reward led to the same interval being perceived longer by human participants (Failing and Theeuwes, 2016; Toren et al., 2020), although one study directly manipulating dopamine signaling in mice found the opposite effect (Soares et al., 2016). However, caution needs to be taken in comparing results on interval perception with those on interval production (Coull et al., 2013), although there is some evidence of shared psychological substrates (Keele et al., 1985; Ivry and Hazeltine, 1995).

Considering that the RewP typically occurs 250-350 ms post-feedback, it is surprising that we did not reliably observe a RewP in the fast tempo which had a target interval of 400 ms. One possibility is that there might be a shift in timing strategies across different tempos. In the slower tempos, participants may rely more on a feedback-based, discrete interval timing system, while the fast tempo may tap into a more automatic and motoric timing system where participants rely more on sampling from their internal interval representation (Lewis and Miall, 2003; Wiener et al., 2011; Petter et al., 2016). Furthermore, feedback processing may interact with tempo speed; the richer the information that the feedback stimuli contain, the smaller the observed RewP amplitude might be (Cockburn and Holroyd, 2018). This study provided directional incorrect feedbacks (early and late) instead of a dichotomous right-or-wrong differentiation, which may require more feedback processing and reduce RewP amplitude in the faster tempos. Overall, this highlights the tradeoff between continuous timing paradigm and RewP amplitude due to a possible shift in timing strategy.

It was argued that the RewP is larger when the feedback is surprising or salient, and that surprise leads to larger behavioral adjustment (Holroyd and Krigolson, 2007; Talmi et al., 2013; but see: Heydari and Holroyd, 2016; Mulligan and Hajcak, 2018). The link between RewP fluctuations and subsequent behavioral adjustment in this study is unlikely to be confounded by surprise about the feedback. This is because the present study used a staircase procedure to ensure that on time feedback always consisted of 50% of trials in each block, eliminating the impact of surprise on the EEG amplitude in response to on time feedback. The different feedback types were also color-coded and randomized across participants to reduce the confound of perceptual salience. This paper adds to the body of literature linking the RewP to subsequent behavioral adjustment, which mostly focused on between-subject level associations across the entire study with a few exceptions (Yasuda et al., 2004; Holroyd and Krigolson, 2007; Cavanagh et al., 2010; Arbel et al., 2013). Several studies that conducted within-subject, trial-based analysis reported non-significant associations between RewP amplitude and timing behavior (Castellar et al., 2010; Cockburn and Holroyd, 2018) . Such results may not be contradictory to our findings. We estimated that each *μV* increase in RewP amplitude slows down the following produced interval by 0.29 ms, equating to a decrease of 0.029% for the 1 s target interval. Given the modest effect size such biasing effect, one possibility is that the limited number of trials from traditional trial-based paradigms (a few hundred trials compared to 1728 trials in this study) may not have the power to detect such effect.

It should be noted that adjustment in timing behavior was not solely dependent on external feedback, but also on internal error monitoring (Miltner et al., 1997; Coles et al., 2001; Ullsperger and Von Cramon, 2003; Danielmeier and Ullsperger, 2011). The neural substrates for internal error monitoring and external reward monitoring are partially separable (de Bruijn et al., 2009). Participants have an internal model of timing which they use to update their belief and modify their behavior (Petter et al., 2016 p.201, 2018). Due to the continuous and ecological property of this paradigm, the internal model may integrate priors about both interval duration and rhythm. First, despite trial-by-trial feedback, participants in this study exhibited a systematic deviation from the target interval. This implies that participants have a prior that biased their produced tempo. Future studies can test this hypothesis by, for example, asking participants to drum with a certain pattern using visual cues without explicit instructions about the speed, and examine whether this natural drumming tempo has an interval below 600 ms. Second, participants’ internal models of rhythm may lead to deviation from target interval as a function of beat location in a pattern (Repp et al., 2011). This study aimed at reducing the influence of internal rhythm by showing participants an explicit drumming pattern, and varying the drumming pattern to be copied. Moreover, the linear mixed model took into considerations where the participant currently was in a pattern, thereby controlling for the influence of rhythm on RT adjustment.

The present study examined how fluctuations in reward processing-related neural activity biases subsequent performance in interval production. We used a continuous drumming paradigm and regression-based analysis to deconvolute overlapping EEG signals. RewP was reliably observed in the slow and medium tempos (target interval 1s or 0.6 s) but diminished in the fast tempo (target interval 0.4s). We found that more positive RewP response to on-time feedback predicts the production of longer interval on the next trial, only in the slow tempo where RewP was the largest. The modest effect size of this behavior-biasing effect of reward highlights the necessity of using a continuous design that allows for more intensive data collection.

## Author Note

This research was funded by the Rhodes Scholarship for China to Yan Yan, a Natural Sciences and Engineering Research Council of Canada (NSERC) Postdoctoral Fellowship to Cameron D. Hassall (PDF 546078 - 2020), a Sir Henry Dale Fellowship from the Royal Society and Wellcome (208789/Z/17/Z) to Laurence T. Hunt., and a NARSAD Young Investigator Award from the Brain and Behavior Research Foundation to Laurence T. Hunt. This research was supported by the NIHR Oxford Health Biomedical Research Centre. The Wellcome Centre for Integrative Neuroimaging was supported by core funding from Wellcome Trust (203139/Z/16/Z).

For the purpose of Open Access, the author has applied a CC BY public copyright license to any Author Accepted Manuscript version arising from this submission.

The authors have no conflict of interest.

## Data Availability Statement

EEG dataset is available at https://openneuro.org/datasets/ds004152/versions/1.1.2. Analysis scripts are available at https://github.com/chassall/drumtrainer.

## Footnotes

1 RewP is also termed Feedback-Related Negativity (FRN) or Error-Related Negativity (ERN) when the contrast is loss minus gain (Miltner et al., 1997; Gehring, 2002; Luu et al., 2003; Proudfit, 2015).

