## Supplementary Materials for "Reward positivity biases interval production in a continuous timing task"

S1 Supplementary results

**Table S1.1** Proportion (and 95% CI) of each feedback type and tempo

| Tempo | feedback | | |
|---|---|---|---|
| | **early** | **late** | **on time** |
| fast | 0.151 (0.116-0.186) | 0.308 (0.273-0.343) | 0.541 (0.534-0.549) |
| medium | 0.301 (0.264-0.339) | 0.168 (0.134-0.202) | 0.531 (0.524-0.537) |
| slow | 0.386 (0.356-0.415) | 0.091 (0.063-0.119) | 0.524 (0.517-0.530) |

**Table S1.2** Full results of the linear mixed model. Model specification: rt_adjustment ~ RewP * tempo * feedback + chunk_location + deviation_from_the_mean * tempo +(1|participant).

| Predictors | Estimates | 95% CI | p |
|---|---|---|---|
| (Intercept) | -0.01077 | -0.01321 – -0.00832 | **<0.001** |
| RewP | 0.00029 | 0.00016 – 0.00042 | **<0.001** |
| tempo [medium] | 0.00705 | 0.00507 – 0.00902 | **<0.001** |
| tempo [fast] | 0.01484 | 0.01286 – 0.01682 | **<0.001** |
| feedback [early] | 0.03417 | 0.03162 – 0.03671 | **<0.001** |
| feedback [late] | -0.03186 | -0.03566 – -0.02806 | **<0.001** |
| chunk location [A1] | -0.00760 | -0.00993 – -0.00528 | **<0.001** |
| chunk location [A2] | -0.00590 | -0.00820 – -0.00360 | **<0.001** |
| chunk location [A3] | 0.00142 | -0.00088 – 0.00372 | 0.227 |
| chunk location [B0] | 0.00853 | 0.00586 – 0.01121 | **<0.001** |
| chunk location [B1] | -0.00681 | -0.00940 – -0.00421 | **<0.001** |
| chunk location [B2] | -0.01203 | -0.01469 – -0.00936 | **<0.001** |

| | | | |
|---|---|---|---|
| chunk location [B3] | -0.00493 | -0.00753 – -0.00233 | **<0.001** |
| chunk location [B4] | -0.00008 | -0.00267 – 0.00251 | 0.951 |
| chunk location [B5] | -0.00490 | -0.00750 – -0.00230 | **<0.001** |
| deviation from the mean | -0.40628 | -0.42620 – -0.38636 | **<0.001** |
| RewP × tempo [medium] | -0.00029 | -0.00047 – -0.00010 | **0.002** |
| RewP × tempo [fast] | -0.00017 | -0.00035 – 0.00002 | 0.076 |
| RewP × feedback [early] | -0.00033 | -0.00053 – -0.00013 | **0.001** |
| RewP × feedback [late] | -0.00035 | -0.00071 – -0.00000 | **0.048** |
| tempo [medium] × feedback [early] | -0.01151 | -0.01512 – -0.00791 | **<0.001** |
| tempo [fast] × feedback [early] | -0.01094 | -0.01492 – -0.00696 | **<0.001** |
| tempo [medium] × feedback [late] | 0.01161 | 0.00671 – 0.01651 | **<0.001** |
| tempo [fast] × feedback [late] | 0.01205 | 0.00747 – 0.01663 | **<0.001** |
| tempo [medium] × deviation from the mean | -0.14825 | -0.18426 – -0.11224 | **<0.001** |
| tempo [fast] × deviation from the mean | -0.20688 | -0.24525 – -0.16851 | **<0.001** |
| (RewP × tempo [medium]) × feedback [early] | 0.00041 | 0.00011 – 0.00070 | **0.007** |
| (RewP × tempo [fast]) × feedback [early] | 0.00023 | -0.00011 – 0.00058 | 0.188 |
| (RewP × tempo [medium]) × feedback [late] | 0.00019 | -0.00025 – 0.00063 | 0.406 |
| (RewP × tempo [fast]) × feedback [late] | 0.00021 | -0.00020 – 0.00062 | 0.319 |
| $N_{participant}$ | 20 | | |
| Observations | 27113 | | |
| Marginal $R^2$ / Conditional $R^2$ | 0.372 / 0.374 | | |

*Note*. Coding of chunk location: A0~A3 were the four chunk locations from pattern A (4/4 time, aaba/aaba/…); B0~B5 were the six chunk locations from pattern B (6/8 time, aaabaa/aaabaa/…); the reference level was set as the first location in pattern A (location A0). CI: confidence interval; RewP: RewP calculated from the regression EEG method.

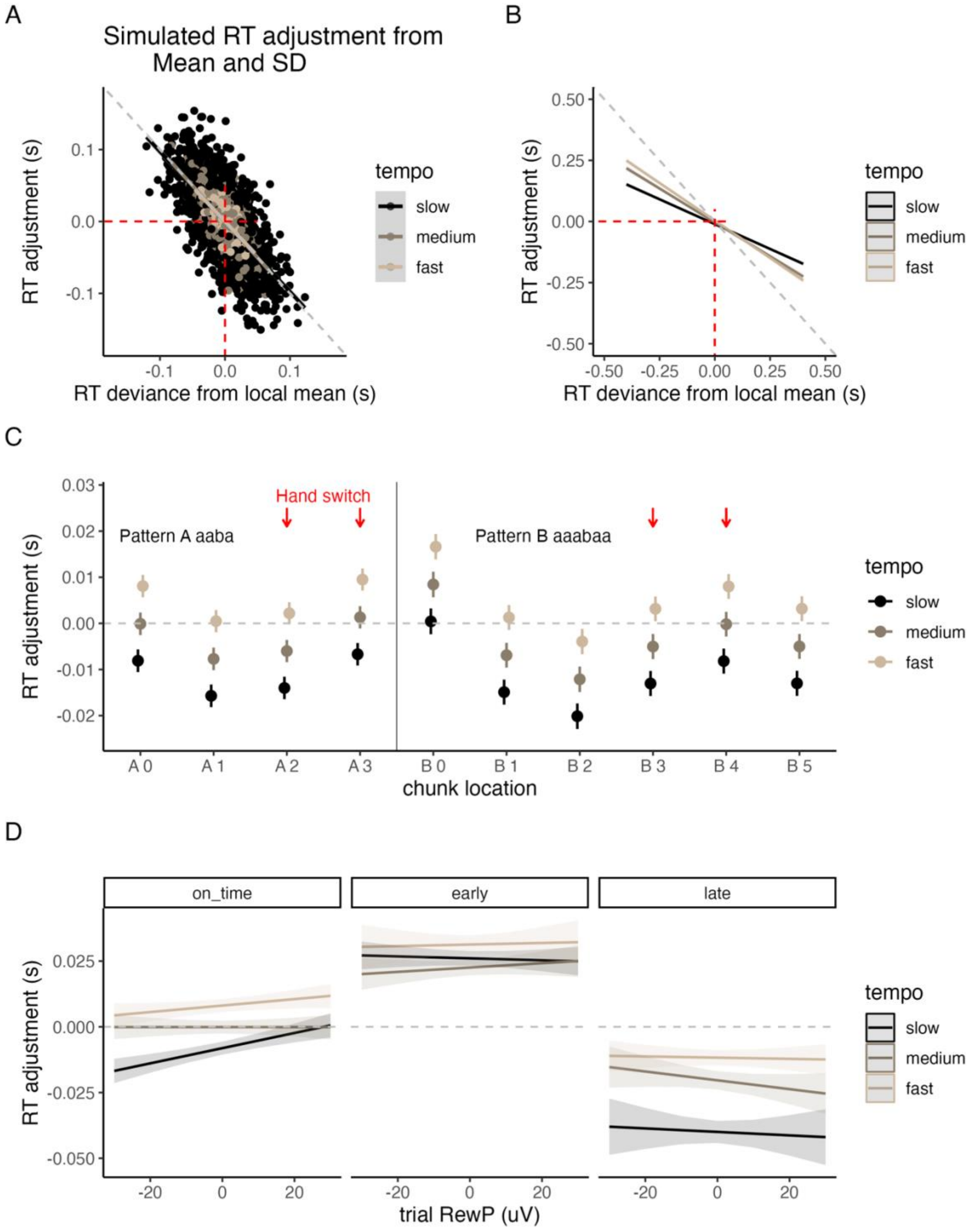


**Figure S1.3** Simulation and the linear mixed model results. (A) Regression to the mean effect in simulated behavioral data. In the simulation, apparent RT adjustment was quantified as the difference between RT in consecutive draws. Deviance from local mean is the difference between last trial's RT and the mean RT of the recent 10 trials. The simulation demonstrated a strong negative association between deviance from the mean and the following apparent RT adjustment. (B) Deviation from local mean contributes to RT adjustment. Larger deviation from local rolling mean in the last trial is associated with RT adjustment on this trial, suggesting regression to the mean. However, the slope of this effect is different from that of (C) Estimates of RT adjustment as a function of chunk location. A0~A3 suggests the four chunk locations in the 4/4 pattern, and B0~B5 suggests the six chunk locations in the 6/8 pattern. (D) The association between RewP amplitude and RT adjustment in different tempos and feedback types. Only the slopes for on-time feedback type in the slow emerged as significant. No slopes in the fast and medium tempo, or in the early and late feedback type were significant.

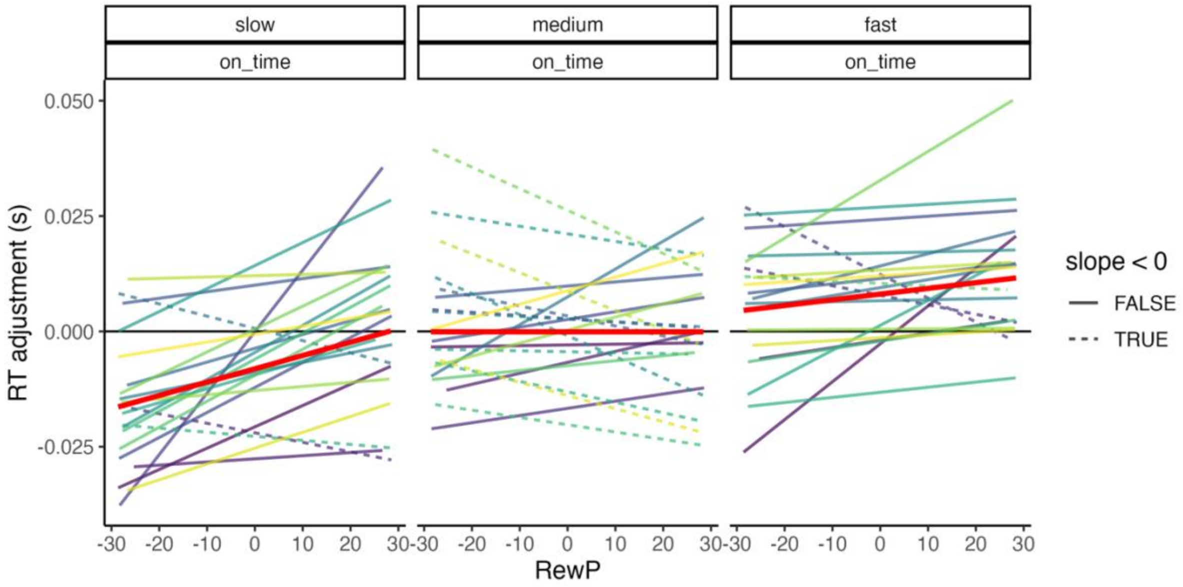


**Figure S1.4** Model estimation for RewP amplitude predicting RT adjustment, using a linear mixed model with variable slope and intercept for each participant respectively. Dashed line indicates slopes that were negative, and the red lines indicate the group-level estimate using the linear mixed model with variable intercept as specified in Table S1.2.

S2 Traditional ERP analysis and its comparison to rERP

*Extraction of traditional Event-related Potentials*

An event-related potential (ERP) is the voltage fluctuation in EEG that occurs as a result of external or internal event (Kappenman and Luck, 2016). Conventional ERP epochs were first created from -200 ms to 600 ms relative to the onset of visual feedback. Epochs were baseline-corrected using the mean voltage from -40 to 0 ms.

Epochs for which the sample-to-sample voltage exceeded 40 μV or the voltage change across the entire epoch exceeded 150 μV were removed.

*RewP amplitude in different tempos*

Two-way within-subject ANOVA (3 tempos × 2 feedback contrasts) on the mean RewP amplitude for each participant suggested a significant main effect of tempo ($F_{(2,38)}$ = 10.223, p < .001, $\eta_p^2$ = 0.350), but not feedback type ($F_{(1,19)}$ = 0.437, p = .517, $\eta_p^2$ = 0.022). There was no significant interaction between tempo and feedback type ($F_{(2,38)}$ = 2.054, p = .142, $\eta_p^2$ = 0.098). Pairwise t-tests with Bonferroni corrections suggested that the RewP amplitude in the slow tempo was significantly larger than in the medium tempo (Mean difference = 1.153, $t_{(1,39)}$ = 3.220, $p_{adjust}$ = .003, Cohen's d = 0.51) and in the fast tempo (Mean difference = 1.464, $t_{(1,39)}$ = 3.423, $p_{adjust}$ = .001, Cohen's d = 0.54); the RewP amplitude in the medium tempo was not significantly different with that in the fast tempo (Mean difference = 0.312, $t_{(1,39)}$ = 1.036, $p_{adjust}$ = 0.307, Cohen's d = 0.16). Post hoc comparisons of the interaction effect revealed that in the medium tempo, the RewP amplitude in response to on time versus early feedback was smaller than in contrast with late feedback (Mean difference = -0.732, $t_{(1,19)}$ = -2.250, $p_{adjust}$ = 0.0365, Cohen's d = -0.50).

**Table S2.1** shows the amplitude of RewP as a function of tempo and feedback contrast type. Noticeably, in the fast tempo, the 95% CI for the early feedback contrast included 0, whereas the 95% CI for the late feedback contrast did not include 0. However, in regression-based analysis, both the early and late feedback contrasts included 0 (see **Table 1** in the main text).

**Table S2.1** Summary statistics of participants' mean RewP amplitude for each tempo and feedback type, derived from traditional ERP analysis.

| tempo | Feedback contrast | mean | SD | 95%CI | | Cohen's d |
|---|---|---|---|---|---|---|
| | | | | lower | upper | |
| fast | On time - early | 0.197 | 2.191 | -0.828 | 1.223 | 0.09 |
| | On time - late | 0.784 | 1.536 | 0.065 | 1.503 | 0.51 |
| medium | On time - early | 0.437 | 0.899 | 0.016 | 0.858 | 0.49 |

| | | | | | | |
|---|---|---|---|---|---|---|
| | On time - late | 1.168 | 1.202 | 0.656 | 1.731 | 0.97 |
| slow | On time - early | 2.286 | 1.634 | 1.522 | 3.051 | 1.40 |
| | On time - late | 1.624 | 2.535 | 0.437 | 2.810 | 0.64 |

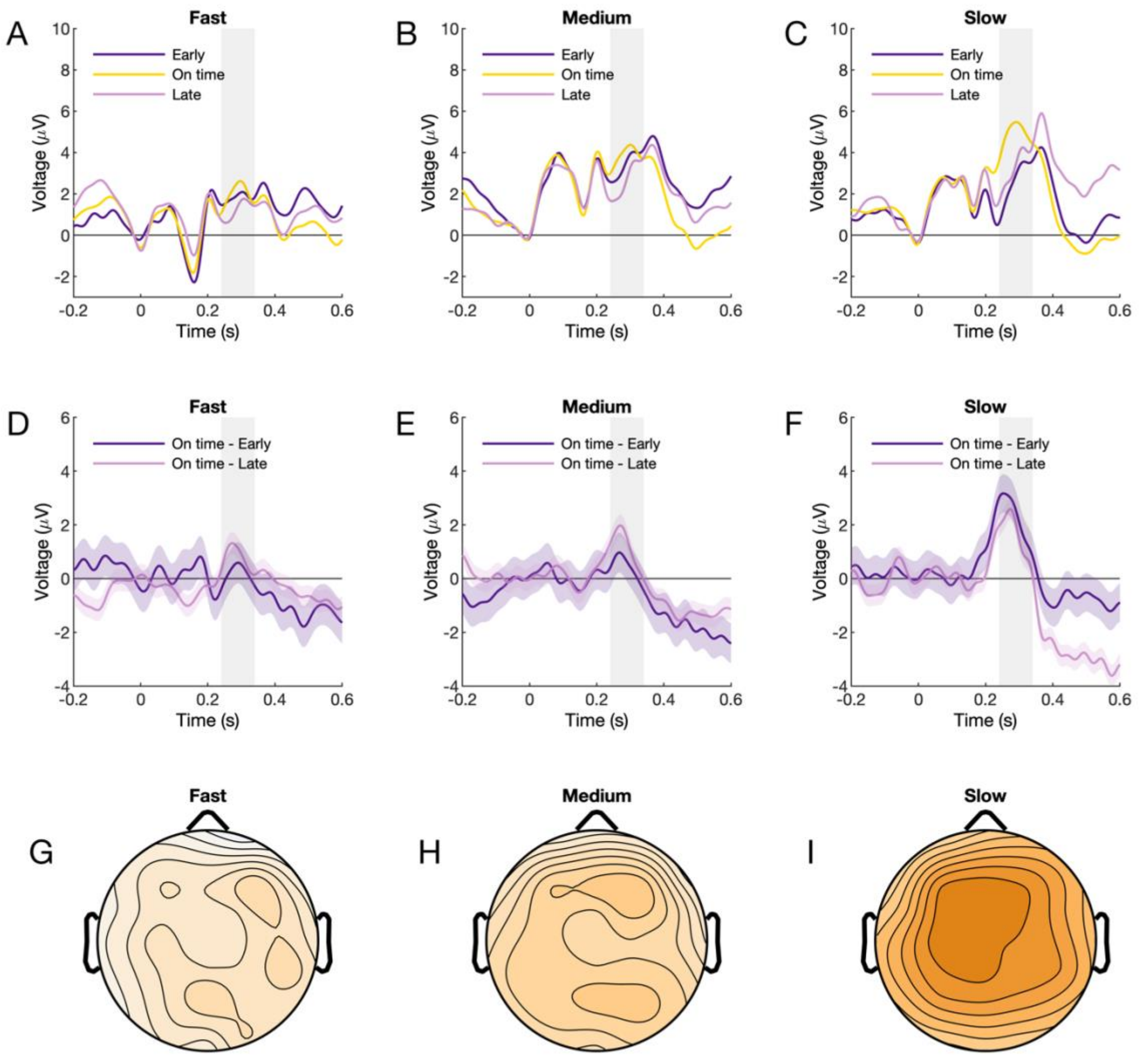


**Figure S2.1** RewP analysis from traditional ERP approach. (A-C) ERP by tempo and feedback type. The RewP time window as reported in Sambrook and Goslin (2015), was highlighted in grey (240-340 ms). (D-F) The RewP wave was calculated as the contrast between correct and incorrect feedback type. RewP amplitude increased as a function of target interval. (G-H) The topography of RewP, averaged between early and late feedback type. In the slow tempo, the peak amplitude was located at electrode FCz.

*Trial-by-trial analysis*

We derived trial-to-trial traditional ERP by taking the mean EEG amplitude within the above-mentioned RewP time window. All other procedures were the same as regression-based EEG analysis. The residual trial-to-trial EEG in the RewP time window derived from regression-based EEG was highly correlated with that derived from raw EEG (random between-subject intercepts; $B = 0.922$, $t = 468.840$, $p < .001$, Cohen's $d = 5.70$). We conducted the same linear mixed model using trial-to-trial EEG extracted from traditional ERP approach. RewP following on time feedback was still a significant predictor of RT adjustment in the slow tempo, although the effect size was smaller, $B = 2.2\times10^{-4}$, $t = 3.378$, $p < .001$, Cohen's $d = 0.041$. RewP did not significantly predict RT adjustment in the medium tempo, $B = -0.08\times10^{-4}$, $t = -1.202$, $p < .001$, Cohen's $d = -0.019$, but significantly predicted RT adjustment in the fast tempo, $B = 2.7\times10^{-4}$, $t = 4.140$, $p < .001$, Cohen's $d = 0.053$.

### S3 Cluster-based permutation test to clarify the time of feedback processing associated with subsequent behavioral adjustment

Having shown that trial-by-trial RewP predicts behavioral adjustment in the next trial, it is still unclear whether the influence of neural response on RT adjustment was limited to the RewP time window, or if it is a general effect spanning across the entire trial duration. To examine this question, we conducted a data-driven analysis to identify the time windows during on-time trials where the EEG activity predict RT adjustment in the following trial. Only on-time trials were selected because the behavior-biasing effect was exclusively identified for on time feedback type. We constructed linear mixed models similar to the one used in **Section 2.5.6**, again using slow tempo and on time feedback type as the reference levels. Instead of the mean amplitude of residual EEG in the RewP time window (240 to 340 ms), these linear mixed models used the residual EEG at FCz for each 4ms time window from -200 ms to 600 ms relative to the onset of the feedback (i.e., -200 to 196 ms, …, 596 to 600 ms). The linear regressions rendered a time series of 200 t value, for each tempo respectively. To identify periods of time where the t values are significant, the t value time series underwent a cluster-based permutation test, with a minimal cluster size of 40 ms (10 time points) and a threshold at $t = 2$. The permutation test was conducted by temporally shuffling the t values in time for $N = 10{,}000$ times, generating a null distribution of the significant cluster size that could arise out of chance. The p value of a cluster is calculated as the proportion of permutations where the permuted cluster size exceeds an observed cluster, and clusters of $p < .05$ were considered as significant. Under the pre-defined threshold, three significant clusters emerged for the t value series in the slow tempo (**Figure S3.1**). The three windows were 20-84 ms ($p = 0.0397$), 256-324 ms ($p = .0.0252$), and 432-556 ms ($p < .0001$). The first and second clusters positively predict RT adjustment, and the third cluster negatively predicted RT adjustment. Notably, the second EEG window between 256 and 324 ms roughly corresponded to the pre-refined time window for RewP (240-340ms). No other significant time windows emerged for fast tempo or medium tempo.

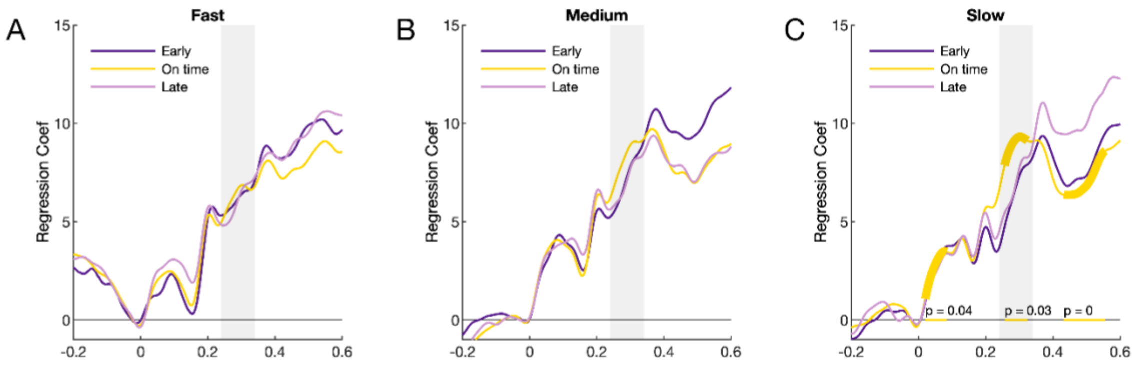


**Figure S3.1** Data-driven exploration of the EEG window that predicts RT adjustment following on time feedback. All p values were unadjusted. The grey area marks the RewP time window as reported in (Sambrook and Goslin, 2015). Three time windows, marked in yellow, were identified as significant clusters with $p < .05$.
